## Supplementary figures and images for "HEB collaborates with TCR signaling to upregulate *Id3* and enable γδT17 cell maturation in the fetal thymus"

### Figure 1 - figure supplement 1

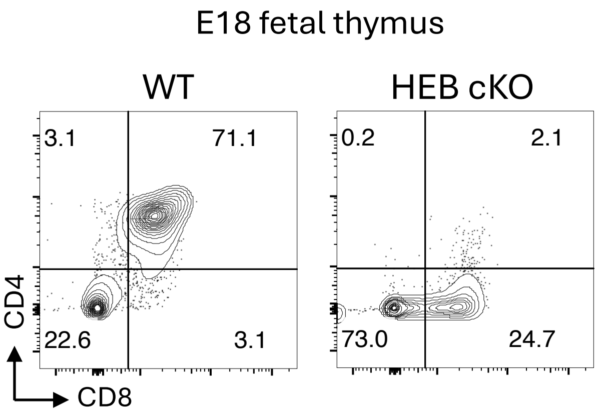

### Figure 2 - figure supplement 2

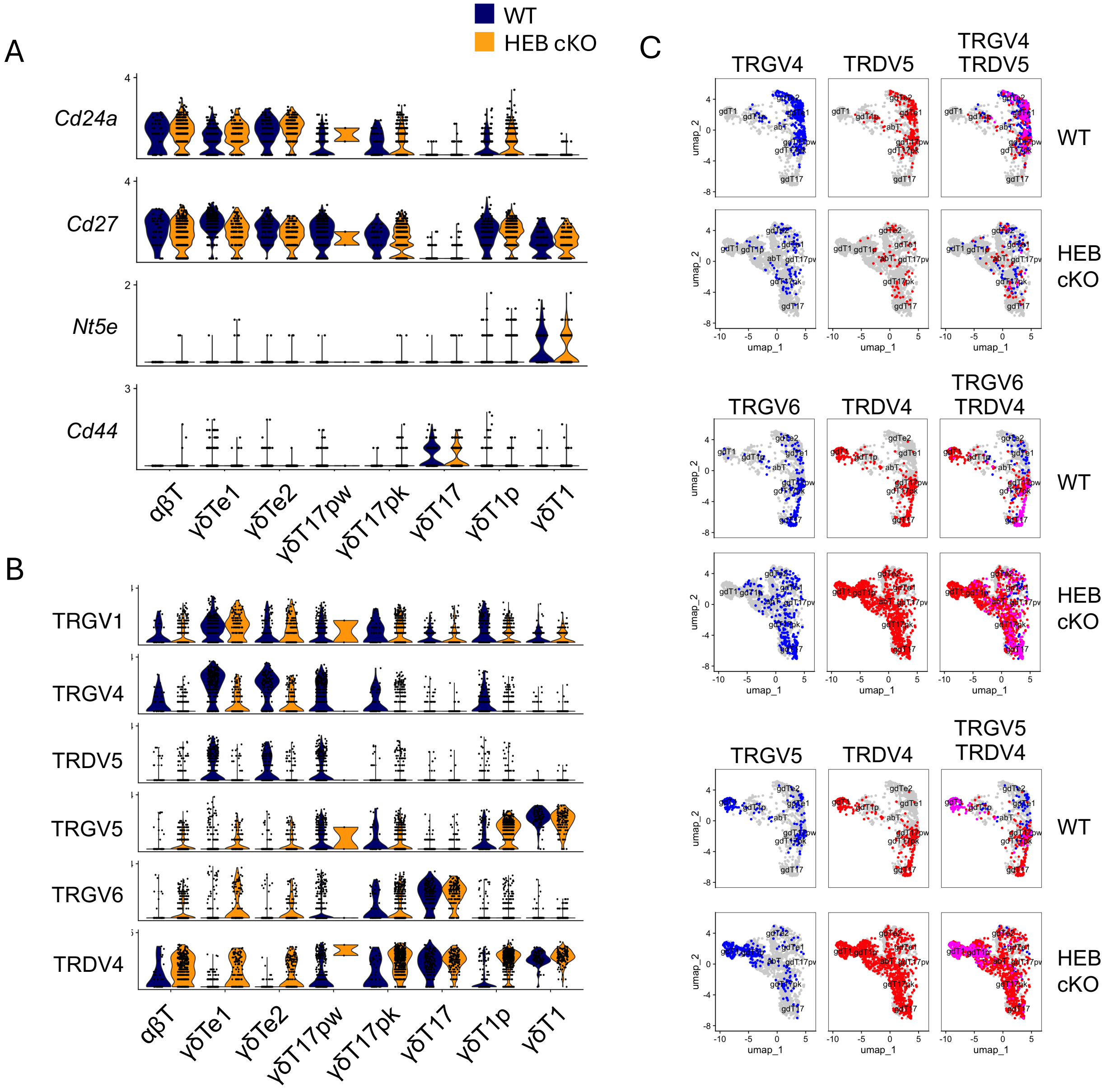

### Figure 4 - figure supplement 3

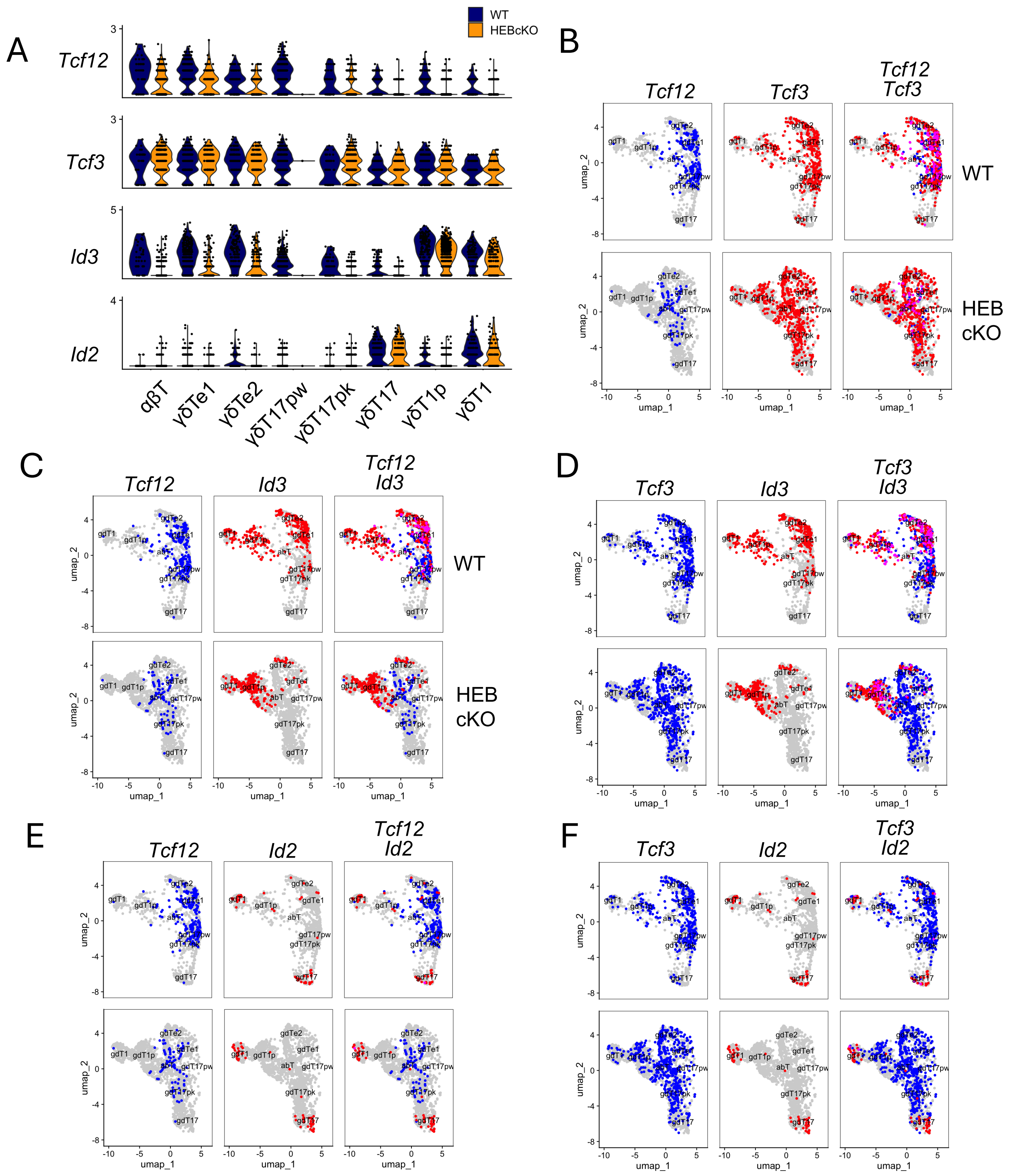

### Figure 5 - figure supplement 1

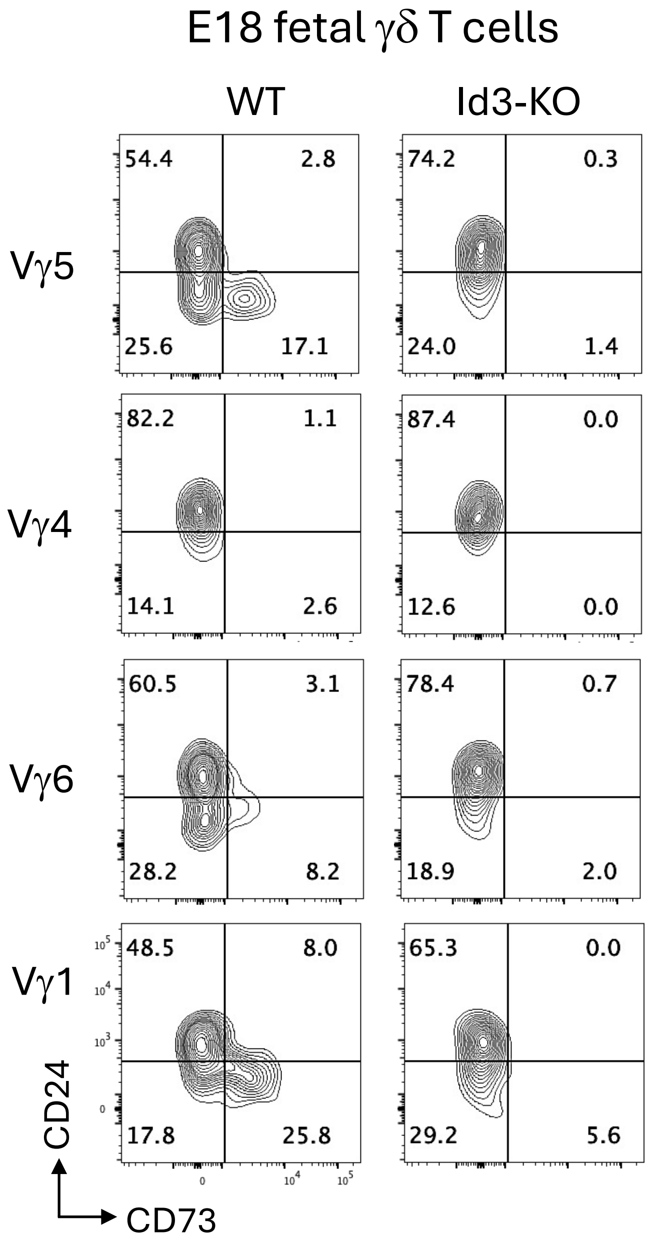

### Figure 6 - figure supplement 1

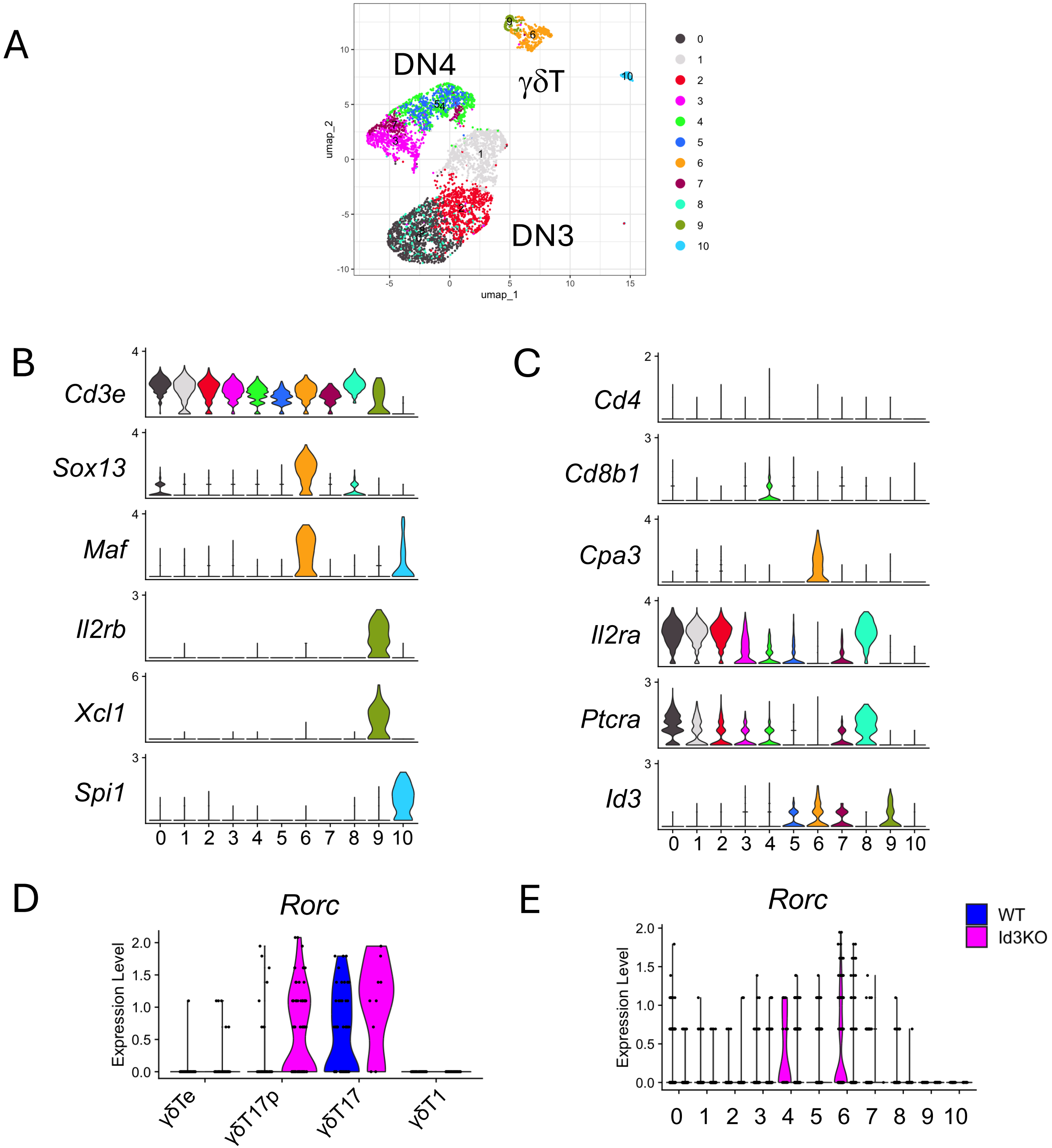

### Figure 6 - figure supplement 2

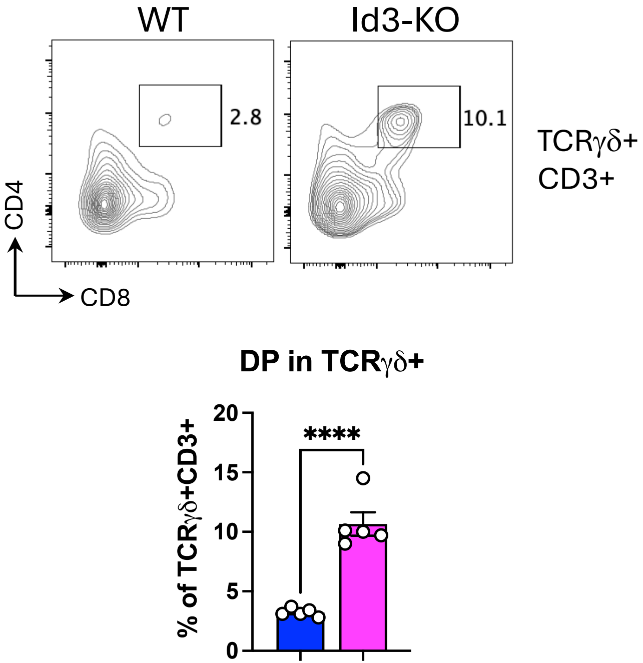

### Figure 7 - figure supplement 1

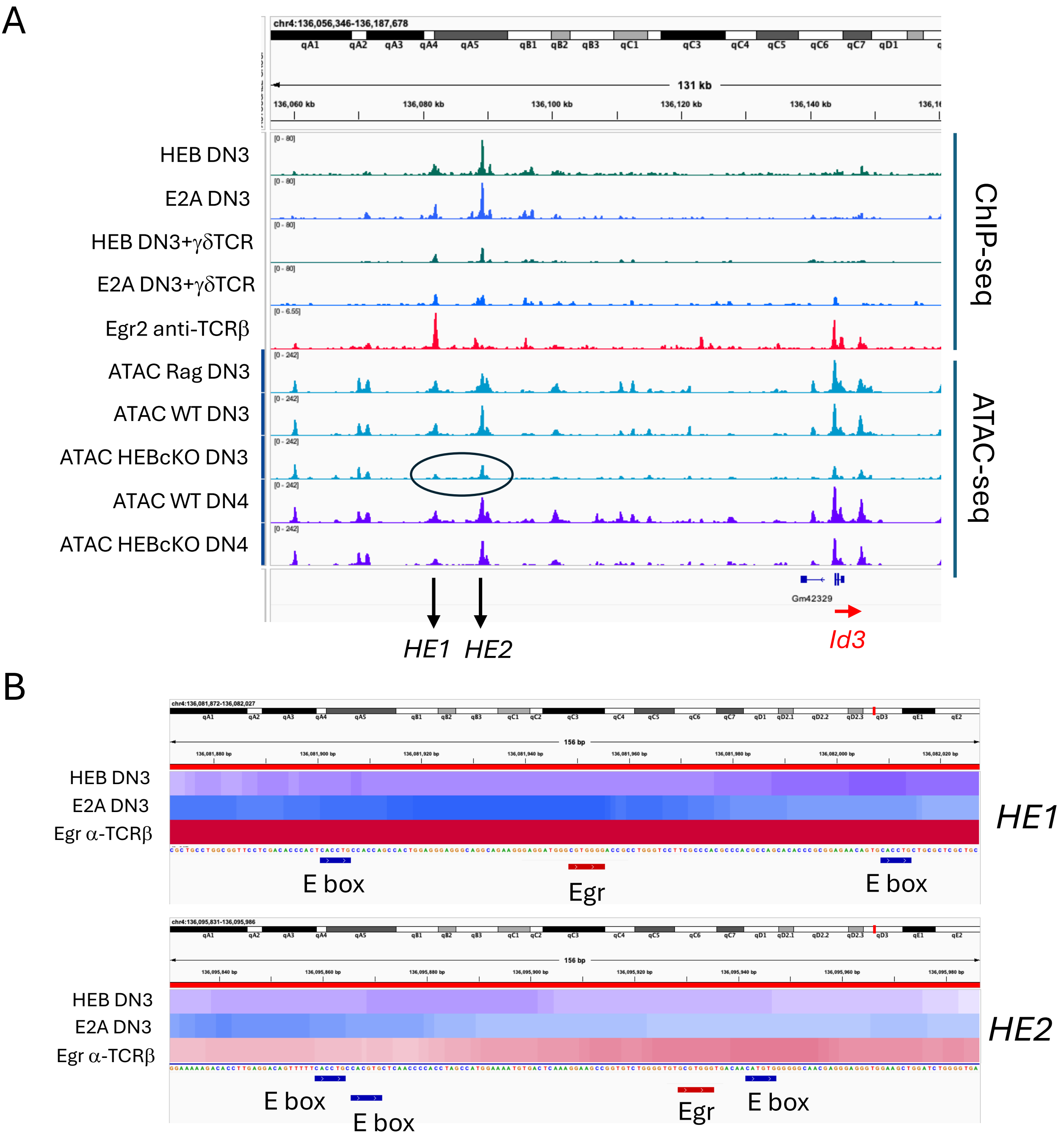

### Figure 7 - figure supplement 2

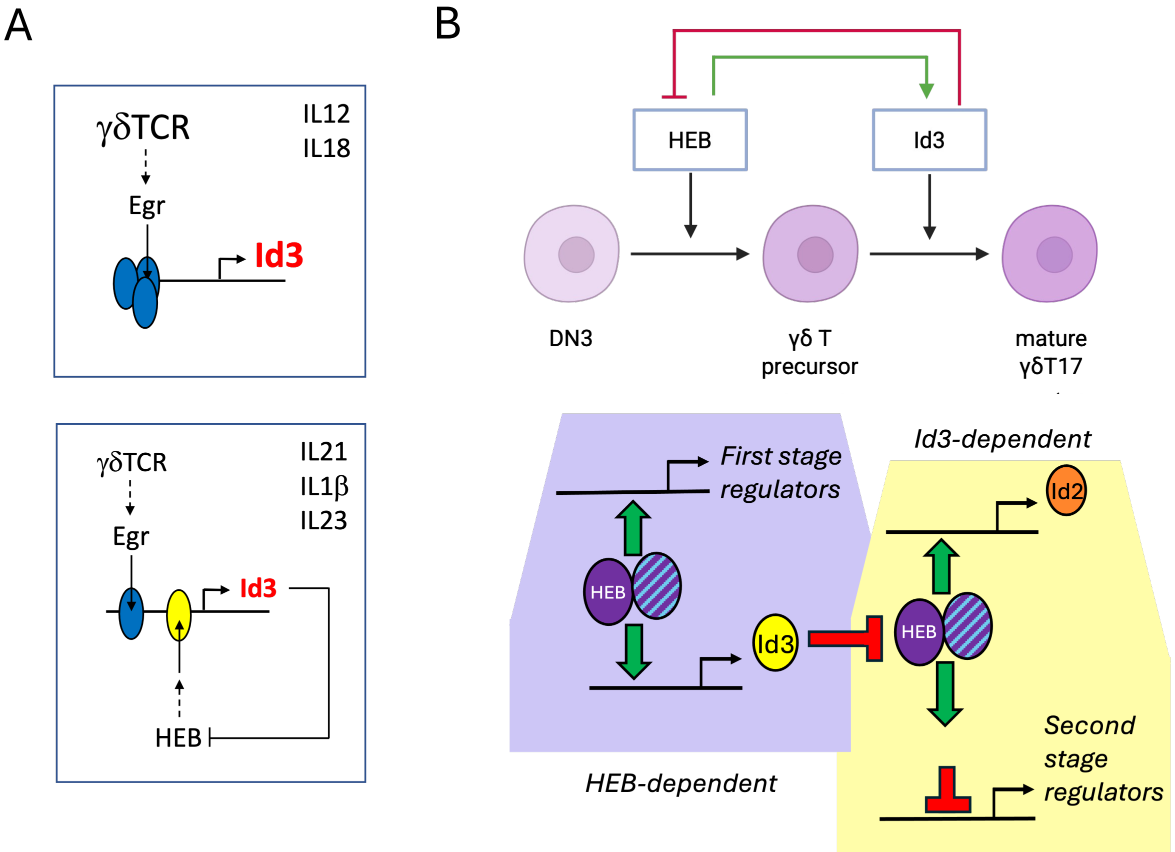
